# Geographic variation in resistance of the invasive *Drosophila suzukii* to parasitism by the biological control agent, *Ganaspis brasiliensis*

**DOI:** 10.1101/2024.01.18.576198

**Authors:** Oscar Istas, Marianna Szűcs

## Abstract

Host-parasitoid interactions are often tied in coevolutionary arms races where parasitoids continuously have to evolve increased virulence as hosts evolve increased resistance. Invasion theories predict that when a host is introduced to a novel region without its coevolved natural enemies, they will evolve lower defenses. Resistance may also differ geographically and temporally due to abiotic and biotic factors. We investigated spatial, temporal and host plant related differences in resistance of the invasive *D. suzukii* in seven geographically distinct populations in Michigan and of one population from Oregon against a coevolved parasitoid, *Ganaspis brasiliensis.* Encapsulation rates (resistance) of *G. brasiliensis* eggs by *D. suzukii* reached 39% in August and 48% in September regionally. These relatively high levels of resistance in North America contrast with expectations and may be due to the low levels of competition *D. suzukii* experiences in the invaded range. Encapsulation rates of *D. suzukii* differed regionally and temporally but not between fruit types. The northernmost site with the lowest encapsulation rate had the highest rate of parasitism suggesting that parasitoids may be able to detect the defensive capacities of their hosts and adjust attack rates accordingly. The lowest encapsulation rates at the northernmost and thus overall coldest site indicate a negative effect of temperature on resistance. However, temporal differences in resistance from August to September were not consistent among sites. These results indicate that there can be regional and temporal variation in the outcome of host-parasitoid interactions between *D. suzukii* and *G. brasiliensis*. This may influence the efficacy and biocontrol potential of *G. brasiliensis* that has been recently approved for field releases against *D. suzukii* in North America.

## Introduction

Insect host-parasitoid systems have provided classic models for research on coevolutionary dynamics (Kraaijeveld et al., 1995; Kraaijeveld et al., 1998; Thompson, 2005; Thompson, 2005). Insect parasitoids lay their eggs inside or on the host and feeding by the developing larvae eventually kills the host. Since only one party survives this interaction, there is strong selection pressure for prey to escape parasitism by mounting physiological and behavioral defenses (resistance) and for parasitoids to overcome host defenses and increase developmental success (virulence) (Godfray 1994). Research focusing on European *Drosophila* species and their native parasitoids shows that there can be regional differences in virulence and resistance (Kraaijeveld et al., 1995; Kraaijeveld & Godfray, 1999). This indicates that it is not solely interactions between parasitoids and hosts that can lead to changes in these important traits, but that the environmental effects on a host-parasitoid community may play a role as well. Both host resistance and parasitoid virulence can also evolve rapidly (Jalvingh et al., 2014; Cavigliasso et al., 2019; Moiroux et al., 2018), and the maintenance of both can be costly (Fellowes et al., 1998; Kraaijeveld & Godfray, 2002; Kraaijeveld & Godray, 1997; Kraaijeveld et al., 2001).

Invasion theories posit that escape from specialist natural enemies (Enemy Release Hypothesis - ERH) (Keane & Crawley, 2002) may allow for the re-allocation of costly defenses towards growth and development (Evolution of Increased Competitive Ability - EICA) (Honor & Colautti, 2020; Blossey & Notzold, 1995). Thus, the combined prediction of ERH and EICA is that invasive species will evolve lower defenses in the introduced range in the absence of specialist parasitoids compared to the native range. Invasive insect pests are often the target of classical biological control programs that focus on the introduction of specialized natural enemies from an invasive species’ native range. As invasive hosts are reunited with their co-evolved natural enemies, reciprocal selection pressures are restored and a new coevolutionary arms race can ensue in the introduced range. The outcome of this new arms race can determine the success or failure of the biocontrol program and thus it is important to understand the mechanisms that influence host-parasitoid interactions in the introduced ranges.

The focus of this study is *Drosophila suzukii* (Diptera: Drosophilidae) that invaded the Americas and Europe in 2008 (Asplen et al., 2015). *Drosophila suzukii* has a wide host range, attacking economically important crops such as raspberries, blueberries, strawberries, and cherries, as well as numerous wild hosts, such as dogwood, pokeweed, choke cherry, and elderberry (Asplen et al., 2013; Poyet et al., 2015; Lee et al., 2019). The extreme polyphagy coupled with high fecundity (> 600 eggs per female) and short generation times (∼14 days at 22°C) make *D. suzukii* difficult to control (Wang et al., 2018). Furthermore, the large population sizes and multiple generations per season also provide ideal conditions for rapid evolution.

*Drosophila suzukii* differs from native North American and European drosophilids in two important traits. Firstly, it has a serrated ovipositor that allows for the piercing of harder, still ripening fruit (Asplen et al. 2005). Therefore, it can occupy a novel niche that is unavailable to other drosophilids that can only lay eggs in overripe, fallen, or rotting fruit (Asplen et al., 2015; Lee et al., 2019). Secondly, *D. suzukii* has a much stronger immune response to parasitism than native drosophilids (Kacsoh & Schlenke, 2012; Poyet et al., 2013). Only two cosmopolitan, generalist pupal parasitoids have been able to develop on *D. suzukii* in the introduced ranges, but their attack rates have remained below 10% leaving *D. suzukii* largely free from parasitism (Chabert et al. 2012; Lee et al. 2019). The eggs and/or larvae of native larval parasitoids are frequently encapsulated and killed by *D. suzukii* as it mounts a cellular and humoral immune response (Poyet et al., 2013; Kacsoh & Schlenke, 2012). *Drosophila suzukii* has a high haemocyte load that is shown to correlate with increased levels of resistance to parasitism (McGonigle et al. 2018, Poyet et al. 2013). However, maintaining high levels of such constitutive resistance is costly and can lead to trade-offs in the form of increased larval mortality and slower development in competitive environments and when food is limited (Fellowes et al., 1998; Kraaijeveld & Godfray, 1997; Kraaijeveld et al., 2001; McGonigle et al., 2017).

Given the scarcity of co-evolved specialist parasitoids and the cost of maintaining high levels of defenses, the expectation is that *D. suzukii* may have evolved lower resistance against parasitism in its introduced ranges compared to the native range. There is little data to confirm such predictions, however, and the one study that has directly compared resistance of introduced and native *D. suzukii* populations showed the opposite pattern. Poyet et al. (2013) found that haemocyte loads in three invasive *D. suzukii* populations from France were twice as high as two native populations from Japan. The higher haemocyte levels correlated with higher encapsulation rates of a European larval parasitoid, *Leptopilina heterotoma* (Poyet et al. 2013).

In this study, we investigated spatial, temporal and host-plant related patterns of resistance of *D. suzukii* populations by sampling eight geographically distinct populations in Michigan and Oregon over a two-month period in three different types of host plants. We assayed resistance using a recently-approved biological control agent, the larval parasitoid *Ganaspis brasiliensis* (Hymenopters: Figitidae) that is a coevolved natural enemy of *D. suzukii* from its native range. We predicted that the North American *D. suzukii* populations will have low resistance to parasitism and that there would be geographic variation in their resistance.

## Materials and Methods

### Parasitoid rearing

A laboratory colony of *G. brasiliensis* was established at Michigan State University (MSU) using 50 adult males and 50 adult females from a USDA-APHIS laboratory in Newark, NJ in February 2022. The *G. brasiliensis* population used in our experiments was established using 15 adult males and 15 adult females from this MSU-based colony in May 2022. To rear *G. brasiliensis*, 150 grams of store bought conventionally grown blueberries were placed into 25 cm x 19 cm x 25 cm plastic containers (PrepNaturals) and sprinkled with 1 teaspoon of instant dry yeast (Fleischmann’s Active Dry Yeast) each to reduce mold buildup and ensure infestation from flies (Rossi-Stacconi et al., 2022). Two containers were placed without lids in a 30 cm x 30 cm x 30 cm mesh cage (RestCloud), and 100 *D. suzukii* flies (50 males and 50 females) were released to infest the blueberries for 48 hours. The *D. suzukii* used to infest the fruit originated from our laboratory colony that was funded in 2018 and augmented with wild-caught individuals each year. The fly rearing proceeded as described in Jarrett et al., (2022) using the standard DSSC cornmeal diet in incubators set at 25±2°C, 70±5% relative humidity, and a 16L:8D photoperiod.

The containers with the fly infested blueberries were then removed from the cages, and 30 *G. brasiliensis* individuals (15 male and 15 female) were added to each to parasitize *D. suzukii* larvae for five days. During parasitism containers were covered with lids that had fine mesh windows for ventilation and a strip of honey underneath for the adult parasitoids to feed on. The parasitoids were removed after 5 days and reused for a next round of parasitization of a new batch of *D. suzukii* infested blueberries. This process was repeated 3-4 times until most adult parasitoids died to maximize population growth. Following parasitization, the plastic containers were held in the same incubators used for fly rearing and were checked twice a week to remove any emerging *D. suzukii*. After approximately 28-35 days *G. brasiliensis* adults emerged, which were then used to start the next parasitoid generation.

### Field sampling of Drosophila suzukii populations

We sampled seven *D. suzukii* populations in Michigan in 2022 following a north to south and east to west grid (Fig. 1) and received samples from one location in Corvallis, Oregon. The Oregon site was included in the sampling to represent a location relatively close to the first detection of *D. suzukii* that occurred in California (Asplen et al. 2015). The distance between the northernmost and southernmost field sites in Michigan was 400 km, and the westernmost and easternmost sites were about 320 km apart. Most sites were conventionally managed mixed orchards, while the site in Oregon and one site in Michigan were organically managed mixed fruit and vegetable farms (Table 1). *Drosophila suzukii* was collected from each location once in August and in September. At each date three types of fruit samples were collected: 1) managed fruit (cherries, raspberries, or blueberries), 2) unmanaged, wild fruit, and 3) for a standardized sample we used banana traps. For the managed fruit collection 0.5 kg of conventionally grown (i.e., pesticide sprayed) fruit was picked from cultivated plants and fallen fruit from the ground. For the unmanaged fruit samples, 0.5 kg of fruit was collected from brambles and other wild soft fruit bearing plants (i.e., blackberries or pokeweed) that were located along field edges. Each fruit sample was collected by hand and from a minimum of three different plants. For the banana traps, one halved banana was placed in a 454 ml lidded red solo cup with several holes drilled into the bottom and lid to allow flies to enter and rainwater to leave. Wires were threaded through holes at the top of the cup and used to hang the traps from trees located along the edge of orchards. At each site three banana traps were placed and left for 48 hours to be infested by fruit flies. An eighth field site located in Corvallis, Oregon was sampled similarly in August and September 2022 and the collected fruit was mailed to MSU to rear out fruit flies and for subsequent resistance assays.

**Figure 1.**
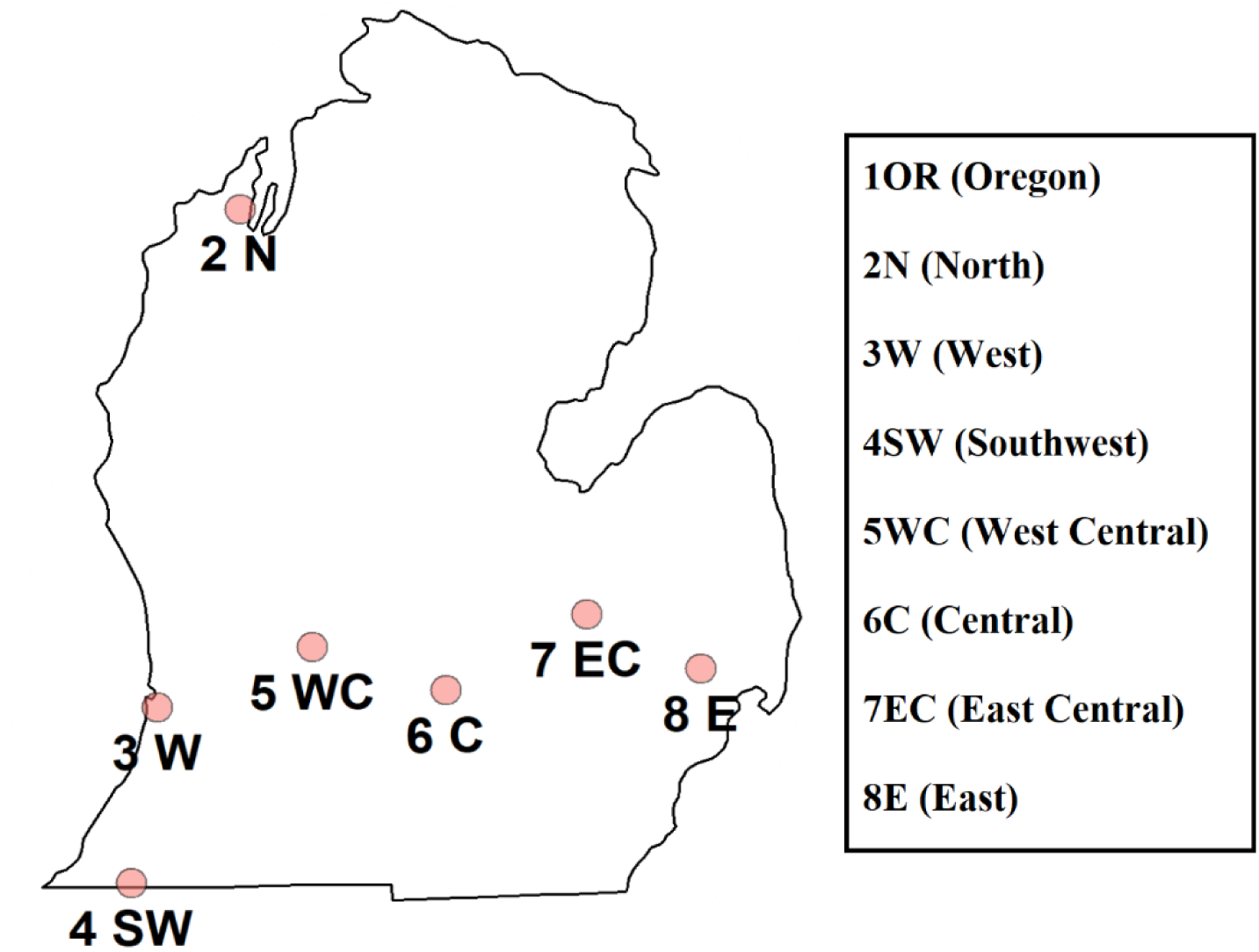
Sites sampled and assayed for *Drosophila suzukii* resistance in 2022. An additional location was sampled in Corvallis, OR. For coordinates, site and sampling information see Table 1.

**Table 1.**
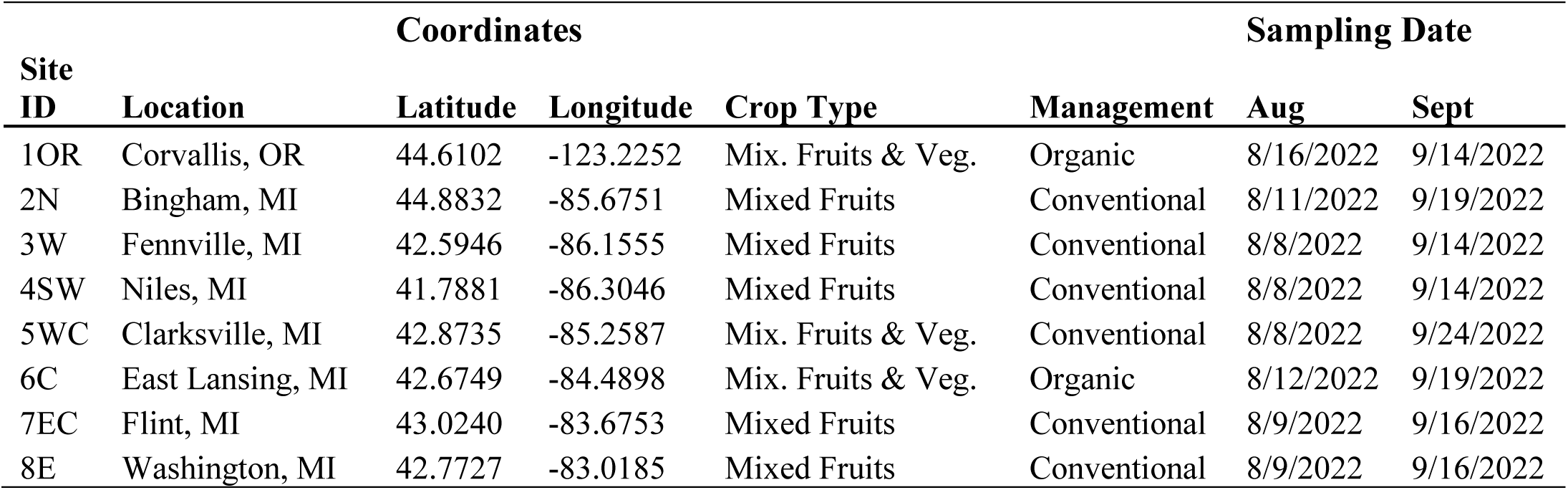
Study locations, site characteristics, and sampling dates. Site ID refers to locations as shown in Figure 1. N = North, W = West, SW = Southwest, WC = West Central, C = Central, EC = East Central, and E = East.

| Site ID | Location | Coordinates |  | Crop Type | Management | Sampling Date |  |
| --- | --- | --- | --- | --- | --- | --- | --- |
|  |  | Latitude | Longitude |  |  | Aug | Sept |
| 1OR | Corvallis, OR | 44.6102 | -123.2252 | Mix. Fruits & Veg. | Organic | 8/16/2022 | 9/14/2022 |
| 2N | Bingham, MI | 44.8832 | -85.6751 | Mixed Fruits | Conventional | 8/11/2022 | 9/19/2022 |
| 3W | Fennville, MI | 42.5946 | -86.1555 | Mixed Fruits | Conventional | 8/8/2022 | 9/14/2022 |
| 4SW | Niles, MI | 41.7881 | -86.3046 | Mixed Fruits | Conventional | 8/8/2022 | 9/14/2022 |
| 5WC | Clarksville, MI | 42.8735 | -85.2587 | Mix. Fruits & Veg. | Conventional | 8/8/2022 | 9/24/2022 |
| 6C | East Lansing, MI | 42.6749 | -84.4898 | Mix. Fruits & Veg. | Organic | 8/12/2022 | 9/19/2022 |
| 7EC | Flint, MI | 43.0240 | -83.6753 | Mixed Fruits | Conventional | 8/9/2022 | 9/16/2022 |
| 8E | Washington, MI | 42.7727 | -83.0185 | Mixed Fruits | Conventional | 8/9/2022 | 9/16/2022 |

### Resistance Assays

Fruit from each site and sample type was placed in separate plastic containers (25 cm x 19 cm x 25 cm, PrepNaturals) in an incubator (25 ± 2°C, 70% RH, 16L:8D) for 10 days. Emerging adult *D. suzukii* were identified based on morphology, separated from other fruit fly species, and transferred to drosophila vials (2.5 x 9.5 cm) (Genesee Scientific, San Diego, CA, USA) containing 6 mL standard DSSC cornmeal diet. Adult flies were allowed to mate and oviposit in the artificial diet for 48 hours. Three days after the adult flies were removed, 50 second instar larvae were extracted from the artificial diet from each sample. The 50 larvae were divided into five replicates of ten larvae each for each location and sample type combination. An additional set of five replicates was created using second instar larvae from our lab-reared colony of *D. suzukii* to test how reusing the same females may have affected parasitism.

The larvae for each replicate were placed atop a halved blueberry to prevent dehydration inside a 60 x 15 mm plastic Petri dish (Falcon, Corning, NY, USA). Each Petri dish then received one female *G. brasiliensis* wasp that had been isolated with a single male in a drosophila vial for 24 hours to mate. The female wasp was left in the Petri dish for four hours to parasitize D*. suzukii* larvae. Because of the limited availability of *G. brasiliensis* females, we had to reuse the same individuals for multiple rounds of parasitism. Individual females were first used to parasitize *D. suzukii* larvae from each of the eight field sites of the five replicates of the banana traps (n = 40 females). The same females were then moved onto the *D. suzukii* samples collected from managed fruit to parasitize for four hours and after that onto the samples from unmanaged fruit for four hours. An additional five parasitoid females were used in three consecutive times to parasitize *D. suzukii* larvae originating from our laboratory colony to test the effect of reusing females on parasitism rates.

The *D. suzukii* larvae were left in the halved blueberries for 72 hours to allow the effects of parasitism and encapsulation to become visible. The larvae were then removed and examined under a microscope (Nikon SMZ1000, 10x-80x magnification) for evidence of parasitization (the presence of parasitoid eggs) and encapsulation of parasitoid eggs that is indicated by the darkening of the egg as it is encased in melanized host tissue and results in the death of the parasitoid. This process allowed us to see the results of our resistance assays more quickly than if we had waited to examine *D. suzukii* adults after they emerged. For most larvae, these signs were visible under a microscope with sufficient light, but if no evidence of parasitism or encapsulation was immediately clear, larvae were dissected with ophthalmological spring scissors (Fine Science Tools). In each replicate, the number of larvae that were parasitized and the number of encapsulated parasitoid eggs were recorded.

### Statistical Analyses

All statistical analyses were performed using R software 4.2.3 (R Core Team, 2013). Generalized linear models (GLM) from the lme4 package were constructed to determine the impact of fixed effects on our dependent variables, while comparison between effects was performed using the emmeans package (Bates et al., 2015; Lenth et al., 2021). Akaike Information Criterion (AIC) was used with the *aictab* function to select the model with the best fit by comparing the simplest model without any interactions to models with all possible two-way and three-way interactions. All post-hoc pairwise comparisons were performed using the *emmeans* package with Tukey-adjustment (Lenth, 2021). The dependent variables (parasitism rate and encapsulation rate) were treated as binomial outcomes since the proportional values were bounded between 0 and 1. Parasitism was calculated for each female by dividing the number of parasitized hosts (n_p_) by the number of hosts offered (n_0_ = 10) to create a parasitism rate (PR): PR = n_p_/n_0_. Encapsulation rate (ER) was calculated by dividing the number of encapsulated eggs in parasitized hosts (n_e_) by the number of parasitized hosts (n_p_): ER = n_e_/n_p_.

First, to test the effect of reusing the same females on parasitism rate and encapsulation rate GLMs were constructed using month (August or September) and the sequence of parasitism (first, second, or third) in the resistance assays as fixed effects. For these resistance assays, *D. suzukii* larvae from our laboratory colony were offered consecutively three times, with the first parasitism event corresponding to tests conducted on field collected samples from banana traps, the second on managed fruit, and the third on unmanaged fruit samples. Parasitism rate and encapsulation rate of *D. suzukii* larvae collected at different locations, in different fruit samples and at different times were compared using GLMs. In these models, location (the eight field sites), sample type (banana trap, managed fruit, and unmanaged fruit), month (August or September) and the interaction between location and month were included as fixed effects. In the model for parasitism rate, the unmanaged fruit samples were removed from analyses because those had been affected by the reuse of parasitoids in the resistance assays.

## Results

### Parasitism rate

Reusing the same females for parasitization had a significant effect on parasitism rate (F_2,27_ = 5.04, p = 0.014) which did not significantly change between months (F_1,26_ = 0.36, p = 0.55). Parasitism rate declined between the first (mean parasitism ± SE; 89% ± 3.1) and third (73% ± 4.4) use of the females (pairwise comparison: p = 0.014) but not between the first and second (80% ± 4.0) use (pairwise comparison: p = 0.19). As an extension of these results from our laboratory ‘control’ population, it is likely that parasitism rate of *D. suzukii* collected from banana traps (tested first) and from unmanaged fruit (tested third) significantly differed due to our experimental procedures. Therefore, data for unmanaged fruit samples were excluded from subsequent parasitism rate analyses.

Parasitism rate of *D. suzukii* by *G. brasiliensis* differed significantly among samples from different locations (F_7,230_ = 6.89, p < 0.001), between months (F_1,229_ = 11.54, p < 0.001), and between different sample types (F_1,237_ = 4.38, P = 0.014). The northernmost site in Michigan (2N in Figure 1) (81% ± 2.29) had the highest parasitism rate that was significantly higher than the westernmost (3W) (62.5% ± 2.81), southwestern (4SW) (65.7% ± 2.75), west central (5WC) (62.4% ± 2.80) and east central (7EC) (66.3% ± 2.76) locations (all pairwise comparisons: p < 0.05). Samples from Oregon (1OR) (75.4% ± 2.49) had higher rates of parasitism than the westernmost and west central Michigan samples (pairwise comparisons: p < 0.05) (Figure 2).

**Figure 2.**
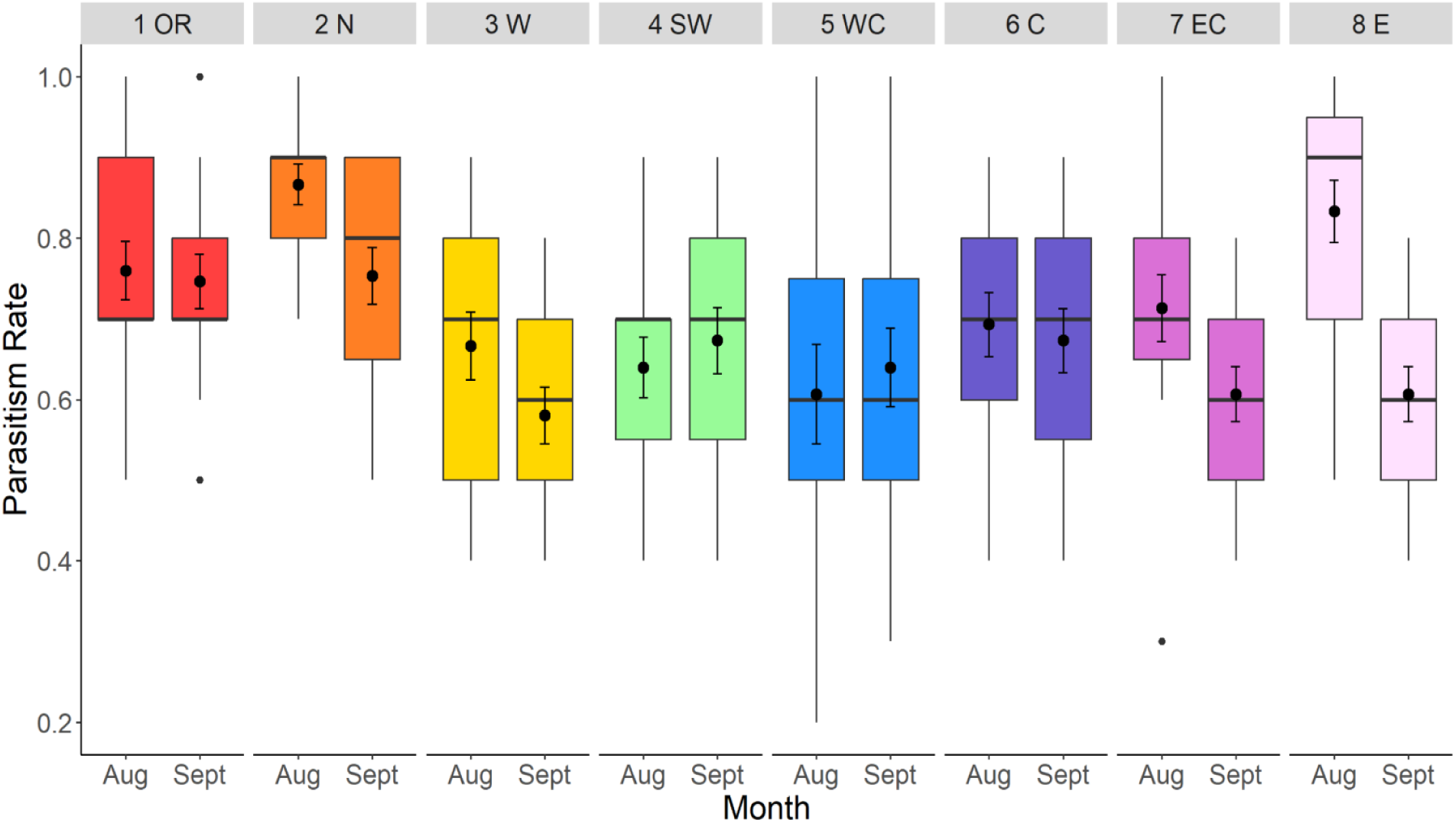
Parasitism rates of *Ganaspis brasiliensis* by eight populations of *D. suzukii* (1 population in Oregon – 1OR and seven populations in Michigan -2N – 8E) sampled in August and September. The numbers on top indicate the sampling locations as shown in Figure 1. Dots indicate outlier observations, the horizontal line indicates the median with the box representing the interquartile range, and vertical lines are 1.5 times the interquartile range. Means and standard errors are shown within each box plot.

Overall, parasitism rates of *D. suzukii* tended to be higher for collections made in August (73.3% ± 1.33) than those collected in September (66.3% ± 1.38) (pairwise comparison: p < 0.001). However, these results were not consistent within locations (month*location interaction: F_7,222_ = 273.96, p = 0.0026). Pairwise comparisons of parasitism rates of August and September samples within sites were only significant at the easternmost site (8E) (p = 0.002) (Figure. 2). The type of sample used to collect *D. suzukii* also led to differences in parasitism rate as larvae collected using banana traps (72.8% ± 1.59) had higher rates of parasitism than those collected from managed fruits (66.1% ± 1.70) (pairwise comparison: p = 0.0105).

### Encapsulation rate

Reusing *G. brasiliensis* females for parasitism did not affect encapsulation rate of parasitoid eggs by *D. suzukii* (F_2,27_ = 3.24, p = 0.055). These results from our laboratory ‘control’ population indicate that reusing females likely did not affect encapsulation rates of *G. brasiliensis* in the field collected samples of *D. suzukii*. Therefore, all three sample types (banana trap, managed and unmanaged fruit) were included in encapsulation analyses of field samples.

Encapsulation rates (resistance) differed among locations where *D. suzukii* was collected from (F_7,230_ = 5.11, p < 0.0001) and between months (F_1,229_ = 10.938, p = 0.0011) but not between fruit sample types (F_2,237_ = 0.19, p = 0.83). The northernmost site that had the highest parasitism rate showed the lowest encapsulation rate (16.1% ± 2.41). Encapsulation rates of the northernmost site were significantly lower than of the Oregon (33.4% ± 3.16), the westernmost (31.1% ± 3.39), and the central Michigan sites (6C) (32.4% ± 3.45) (all pairwise comparisons: p < 0.05) (Fig. 3). The month of sampling affected resistance levels with *D. suzukii* collected in September (29.9% ± 1.66) demonstrating higher encapsulation rates than the August samples (22.3% ± 1.46) (pairwise comparison: p = 0.0006). However, the higher encapsulation rates later in the season were not consistent across sites (location*month interaction: F_7,222_ = 3.868, p = 0.0005). Of the eight sites only the central Michigan location showed significantly higher encapsulation rates in the September samples compared to the August samples (pairwise comparison: p < 0.0001) (Fig. 3).

**Figure 3.**
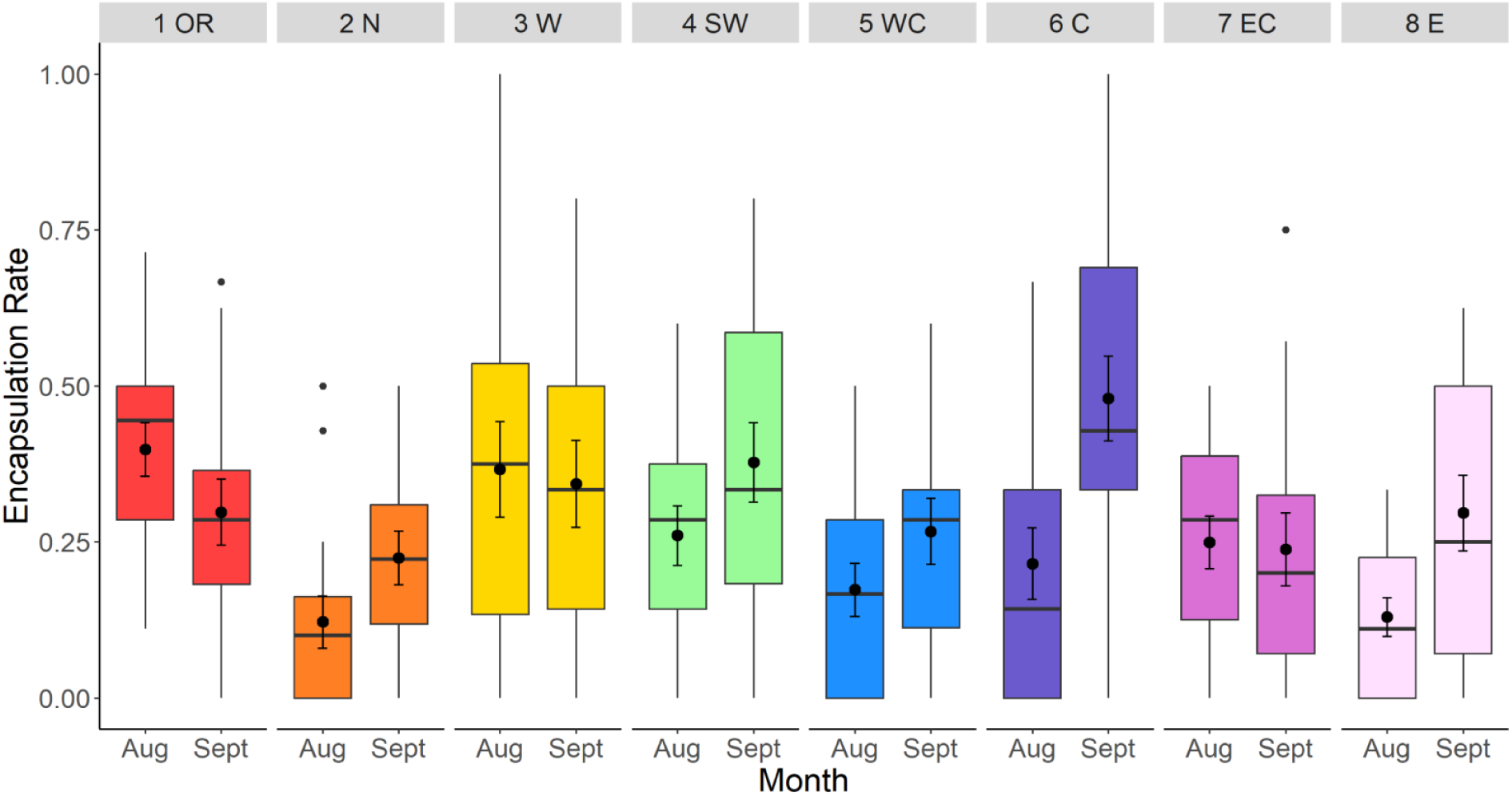
Encapsulation rates of *G. brasiliensis* by eight populations of *D. suzukii* (1 population in Oregon – 1OR and seven populations in Michigan - 2N – 8E) sampled in August and September. The numbers on top indicate the sampling locations as shown in Figure 1. Dots indicate outlier observations, the horizontal line indicates the median with the box representing the interquartile range, and vertical lines are 1.5 times the interquartile range. Means and standard errors are shown within each box plot.

## Discussion

We found geographic and temporal differences in the levels of resistance of invasive *D. suzukii* populations in North America against the specialist larval parasitoid *G. brasiliensis*. In contrast to expectations, resistance of North American *D. suzukii* populations were relatively high. These results can have implications for the incipient classical biological control program that uses mass releases of *G. brasiliensis* and may cause spatial and temporal variation in control success.

Encapsulation rates of *G. brasiliensis* eggs by *D. suzukii* ranged between 12 - 39% in August and 22 - 48% in September regionally. Direct comparison of these results with other studies is difficult because of the different methods used to assess encapsulation. We dissected parasitized larvae to assess parasitism and encapsulation rates (Kacsoh & Schlenke, 2012). The two other studies that assessed the outcome of parasitization by *G. brasiliensis* examined the presence of black capsules in adult flies and found around 5% (Daane et al., 2021) and 15% of encapsulation rates by *D. suzukii* (Girod et al., 2018). These studies also used multiple *G. brasiliensis* populations that had differing host specificity, which can influence parasitism success (Daane et al., 2021; Girod et al., 2018). Even though the encapsulation rate assessments in this study do not account for mortality that could occur during development, the measured encapsulation rates are still relatively high considering that we used the most host specific strain of *G. brasiliensis* (G1) that coevolved with *D. suzukii* in its native range and are used for field releases across the USA (Daane et al., 2021; Hougardy et al., 2022).

Coevolved, host specific parasitoids should have adaptations to effectively overcome host defenses as was shown for the larval parasitoid *Asobara japonica* (Hymenoptera: Braconidae) that only had 6-26% encapsulation rates by multiple strains of *D. suzukii* (Poyet et al., 2013). In contrast, the European parasitoid *L. heterotoma* that did not have a coevolutionary history with *D. suzukii* was encapsulated at rates of 59-87% (Poyet et al., 2013). The strength of resistance correlated with hemocyte levels in the flies that were two times higher in the introduced compared to the native population of *D. suzukii* (Poyet et al., 2013). Thus, both the current study and the one by Poyet et al. (2013) indicate that introduced populations of *D. suzukii* have maintained relatively high levels of defenses against parasitism in contrast to the combined predictions of the ERH and EICA hypotheses (Keane & Crawley, 2002; Honor & Colautti, 2020; Blossey & Notzold, 1995). There is evidence that the maintenance of high haemocyte load is only costly in competitive environments. For example, when food is plentiful, there was no cost to resistance in three *Drosophila* species (McGonigle 2017). It is possible that *D. suzukii* can maintain high levels of defensive compounds without trade-offs because it currently experiences relatively low competition and high resource availability due to its ability to attack still ripening fruit with its serrated ovipositor (Poyet et al. 2013). In contrast, native drosophilids in Europe and North America are restricted to attacking overripe and fallen fruit and thus occupy different niches than *D. suzukii* (Asplen et al. 2015, Lee et al. 2019).

Encapsulation rates varied regionally across Michigan and Oregon. Similar variation in parasitoid resistance has been seen in other *Drosophila* species which have demonstrated population-level differences in encapsulation rates based on geographic location (Kraaijeveld & Van Alphen, 1995; Kraaijeveld & Godfray, 1999; Dubuffet et al., 2007). For example, *D. melanogaster* resistance to one of its larval parasitoids, *A. tabida,* is highest in central Europe (40-60% encapsulation rates) and lower in northern and southern Europe (Kraaijeveld and Godfray 1999). Seven geographically distinct populations of *D. yakuba*, a fly species native to Africa, also show varying encapsulation rates of the larval parasitoid *L. boulardi* that range between 6% and nearly 98% regionally (Dubuffet et al., 2007). Geographic structure in resistance can arise because of spatial and temporal differences in the strength of reciprocal selection between hosts and parasitoids, differences in the genetic variation available in local populations to respond to selection, differences in the wider host-parasitoid community, in abiotic conditions, and how the cost and benefits of resistance are balanced locally in terms of fitness (Kraaijeveld & Godfray, 1999; Kraaijeveld & Van Alphen 1995; Dubuffet et al., 2007). In native host-parasitoid communities where multiple parasitoid and host species interact, the regional difference in community structure can be the most important driver of geographic differences in host defenses (Kraaijeveld & Godfray, 1999). However, in the introduced ranges, host-parasitoid communities can be diminished, as is the case with *D. suzukii* that has experienced very low parasitism rates by native parasitoids over the last decade in North America (Lee et al., 2019).

In the absence of widespread, high-density populations of specialist coadapted parasitoids in the landscape abiotic factors may be the primary factors influencing resistance levels of *D. suzukii* populations. Our results provide some evidence for that since the northernmost site in Michigan had the lowest encapsulation rate. This site is the coldest as it is located over 300 kilometers to the north of other sites in Michigan and lower temperatures have been associated with lower encapsulation ability in insect host-parasitoid interactions (Blumberg, 1991; Fellowes et al., 1999). There were also temporal differences in resistance of *D. suzukii* populations that show the opposite pattern, that is, somewhat increasing encapsulation rates later in the season. However, this difference was only significant between August and September for the central Michigan site (6C in Fig. 1) and *D. suzukii* collected in Oregon showed the reverse pattern of lower encapsulation in September than in August. Thus, it is likely that temperature effects on host resistance are not linear and are mediated by complex interactions between the host, their parasitoids, and the environment (Thomas & Blanford, 2003).

Parasitism rates of *D. suzukii* populations also differed regionally and over time, however, parasitism tended to show opposite patterns than encapsulation. For example, the northern site that demonstrated the lowest encapsulation rates showed the highest rates of parasitism. Similarly, the easternmost site that had low encapsulation rates had relatively high parasitism rates. While encapsulation rate increased from August to September, parasitism rate of the same populations decreased between those months. These patterns suggest that *G. brasiliensis* may be able to assess the resistance ability of larvae and lays fewer eggs in better defended individuals. Such correlations have been shown with the parasitoid *A. tabida* that tended to reject *D. melanogaster* larvae that had high resistance to parasitism (Kraaijeveld et al., 1995). Similarly, in 16 different Lepidopteran species caterpillars with the highest levels of resistance had the lowest levels of parasitism (Smilanich et al., 2009).

The above results indicate that there can be differences in the regional outcome of host-parasitoid interactions that may influence the efficacy of *G. brasiliensis* as a biological control agent of *D. suzukii*. At locations where *D. suzukii* have relatively low resistance the impact of biocontrol may be larger initially, while increased or geographically variable levels of resistance could render biocontrol less successful or more variable regionally. Our results could help to assess the mechanisms that may underlie biocontrol success or failure, and different management approaches may be recommended based on spatial differences in resistance of *D. suzukii* populations. For example, higher parasitoid releases may be necessary at locations that show relatively high levels of *D. suzukii* resistance to increase parasitoid pressure and to ensure that *G. brasiliensis* populations can maintain high densities despite their lower success of development. The higher population sizes and densities could help *G. brasiliensis* to maintain genetic diversity and adaptive potential and to evolve higher virulence over time. An alternative approach may be to release *D. suzukii* flies with low levels of resistance at sites that showed high encapsulation rates to reduce overall resistance by mixing populations. In any case, the baseline data collected here will be valuable to further explore the eco-evolutionary dynamics of host-parasitoid interactions following the release of *G. brasiliensis* across Michigan.

## Data Archiving Statement

Data for this study are available at: *to be completed after manuscript is accepted for publication*.

## Acknowledgements

We are grateful to A. Greenhalgh and J. Lee from Oregon State University for sending us infested fruit from Oregon. We thank Sarah Fitzpatrick, Rufus Isaacs, and Henry Chung for comments on an earlier version of the manuscript. This work was supported by Michigan State University AgBioResearch Project GREEEN (awards: GR21-078 and GR22-064). M. S. was supported by the United States Department of Agriculture National Institute of Food and Agriculture (USDA NIFA) Hatch projects 1017601 and 1018568.

